# Comparison of Two Illumina Whole Transcriptome RNA Sequencing Library Preparation Methods Using Human FFPE Specimens

**DOI:** 10.1101/2021.01.25.428060

**Authors:** Danyi Wang, P. Alexander Rolfe, Dorothee Foernzler, Dennis O’Rourke, Sheng Zhao, Juergen Scheuenpflug, Zheng Feng

**Author notes:** These authors contributed equally to this work (Danyi Wang, P. Alexander Rolfe). Corresponding author, (ZF).

## Abstract

RNA extraction and library preparation from formalin-fixed, paraffin-embedded (FFPE) samples are crucial pre-analytical steps towards achieving optimal downstream RNA Sequencing (RNASeq) results. We assessed the Illumina TruSeq Stranded Total RNA library preparation method and the Illumina TruSeq RNA Access library preparation method for RNA-Seq analysis using 25 FFPE samples from human cancer indications (NSCLC, CRC, RC, BC and HCC) at two independent vendors. These FFPE samples covered a wide range of sample storage durations (3-25 years-old), sample qualities, and specimen types (resection vs. core needle biopsy). Our data showed that TruSeq RNA Access libraries yield over 80% exonic reads across different quality samples, indicating higher selectivity of the exome pull down by the capture approach compared to the random priming of the TruSeq Stranded Total kit. The overall QC data for FFPE RNA extraction, library preparation, and sequencing generated by the two vendors are comparable, and downstream gene expression quantification results show high concordance as well. With the TruSeq Stranded Total kit, the average Spearman correlation between vendors was 0.87 and the average Pearson correlation was 0.76. With the TruSeq RNA Access kit, the average Spearman correlation between vendors was 0.89 and the average Pearson correlation was 0.73. Interestingly, examination of the cross-vendor correlations compared to various common QC statistics suggested that library concentration is better correlated with consistency between vendors than is the RNA quantity. Our analyses provide evidence to guide selection of sequencing methods for FFPE samples in which the sample quality may be severely compromised.

## Introduction

High analytical sensitivity and broad dynamic range render RNA sequencing (RNA-Seq) very appealing for mRNA expression analyses in clinical biomarker development and identification for enabling precision oncology [1]. However, the reliability and accuracy of RNA-Seq data is largely dependent on template RNA quality and input amount as well as the cDNA library preparation methods applied, especially in samples with suboptimal quality that is extracted from FFPE specimens [2]. Several next generation sequencing (NGS) protocols are currently available for the profiling of suboptimal RNA samples, including RNase H, Ribo-Zero, DSN-lite, NuGEN, SMART, and exome capture, each with its own strengths and weakness [3–5].

Among these NGS protocols, Illumina offers two library preparation methods for samples with suboptimal quality: the TruSeq RNA Access library preparation method is based on RNA capture by targeting known exons with exon capture probes to enrich for coding RNAs [4]; the TruSeq Stranded Total RNA library kit with Ribo-Zero rRNA removal (TruSeq Stranded Total RNA) is a method that reduces the highly abundant ribosomal RNAs from total RNA samples using ribosomal capture probes [3]. The performance of the TruSeq Stranded Total RNA and TruSeq RNA Access library preparation kits has been evaluated on well-established human reference RNA samples from the Microarray/Sequencing Quality Control consortium (MAQC/SEQC) [6]. The RNA Access protocol is not only suitable for the profiling of samples of severely compromised quality, but also appropriate for very heterogeneous RNA samples including a wider range of low quantity and extremely low-quality samples [7]. It is essential to conduct a systemic comparison of these protocols using human samples across various cancer indications using different vendors to ensure clinical translatability.

In this study, we compared the performance of the Illumina TruSeq Stranded Total RNA and Illumina TruSeq RNA Access library preparation kits using 25 FFPE samples from patients with five cancers of various sample quality, age of samples, and sample type and between two vendors (Vendor A and Vendor B).

## Materials and Methods

### Clinical samples

Twenty-five FFPE samples from five indications (non-small cell lung cancer (NSCLC), colorectal cancer (CRC), renal carcinoma (RC), breast cancer (BC) and hepatocellular carcinoma (HCC)) of various sample quality, ages of samples (collection year: 1993-2015) and sample type (22 resection vs. 3 core needle biopsy) were procured from three suppliers. The same set of samples were processed using the same protocols from RNA extraction to sequencing and were evaluated using both TruSeq Stranded Total RNA and TruSeq RNA Access library preparation kits at two different vendors (Vendor A and Vendor B).

### RNA extraction and assessment of quality

The RNA extraction of FFPE tumor specimens was performed on five 5μm-deep tissue cuts using the Qiagen RNeasy Mini Kit (Qiagen), according to the manufacturer’s recommendations. Total RNA concentration was measured using Qubit^®^ RNA HS Assay Kit on a Qubit^®^ 2.0 Fluorometer (Thermo Fisher Scientific Inc., Waltham, MA, USA). Integrity was assessed using Agilent RNA 6000 Nano Kit on a 2100 Bioanalyzer instrument (Agilent Technologies, Santa Clara, CA, USA). The RIN score and the percentages of fragments larger than 200 nucleotides (DV_200_) were calculated. According to Agilent 2100 bioanalyzer system assessment and Illumina library preparation input recommendation, the degraded RNA samples can be classified according to their size distribution DV_200_. FFPE RNA samples with DV_200_ >70% are high quality samples, 50-70% are medium quality samples, 30-50% is defined as low quality FFPE while DV_200_ <30% indicates the FFPE RNA is likely too degraded for RNASeq.

### RNA library construction and sequencing

Ribosomal RNA depleted strand-specific RNA libraries were generated with the TruSeq Stranded Total RNA sample preparation kit with Ribo-Zero Gold (#RS-122-2301and #RS-122-2302, Illumina) and transcriptome capture based libraries were generated with the TruSeq RNA Access Library Prep Kit (#RS-301-2001, Illumina). All protocols were performed following the manufacturer’s instructions. Each library was sequenced on an Illumina HiSeq 2500 (Illumina, Inc. San Diego, CA, USA) using V3 chemistry, in paired-end mode with a read length of 2×50bp. Each library was normalized to 20 pM and subjected to cluster and pair read sequencing was performed for 50 cycles on a HiSeq2500 instrument, according to the manufacturer’s instructions. Image analysis, base calling and base quality scoring of the run were processed on the HiSeq instrument by Real Time Analysis (RTA 1.17.21.3) and followed by generation of FASTQ sequence files by CASAVA 1.8 (Illumina, Inc. San Diego, CA, USA). Data are available in the repository NCBI Sequence Read Archive, accession number PRJNA660476.

### Data Processing

All raw data of the samples were processed through a standard RNASeq pipeline to produce counts and transcripts per million (TPM) for each gene in each sample. Reads were aligned with STAR version 2.5.2b [8] against hg19 and the gencode gene annotations version 24 [9]. RSEM version 1.2.29 was then used to quantify and compute TPMs [10]. All downstream processing of the TPM and counts was performed in R version 3.6.1 [11]. QC was performed on the STAR output using Picard (http://broadinstitute.github.io/picard). Unless otherwise noted, we present output only using the fifteen samples which passed QC for all four attempts (two vendors times two protocols). We quantified data quality for a sample in terms of the number of genes detected (count greater than zero) and by the 90^th^ percentile count in a sample (low RNA input often yields extremely high amplification of a few highly expressed genes and few reads at the vast majority of genes, leading to a low 90^th^ percentile count). We assessed the results in terms of spearman and pearson correlation of TPM values between vendors for the same kit, and between the two kits at the same vendor. Spearman correlation measures whether two sets of values are in the same order, even if the relationship is non-linear, while the pearson correlation measures whether the relationship between two datasets is linear. For examples, if the values in one dataset are the square of the values in the other dataset, they would have a high spearman correlation but low pearson correlation.

## Results

### RNA and Library QC Measurements

To evaluate the performance of RNA-seq methods in profiling FFPE samples, we conducted a technical assessment of the two different RNA library preparation protocols on 25 FFPE samples (Fig 1).

**Figure 1.**
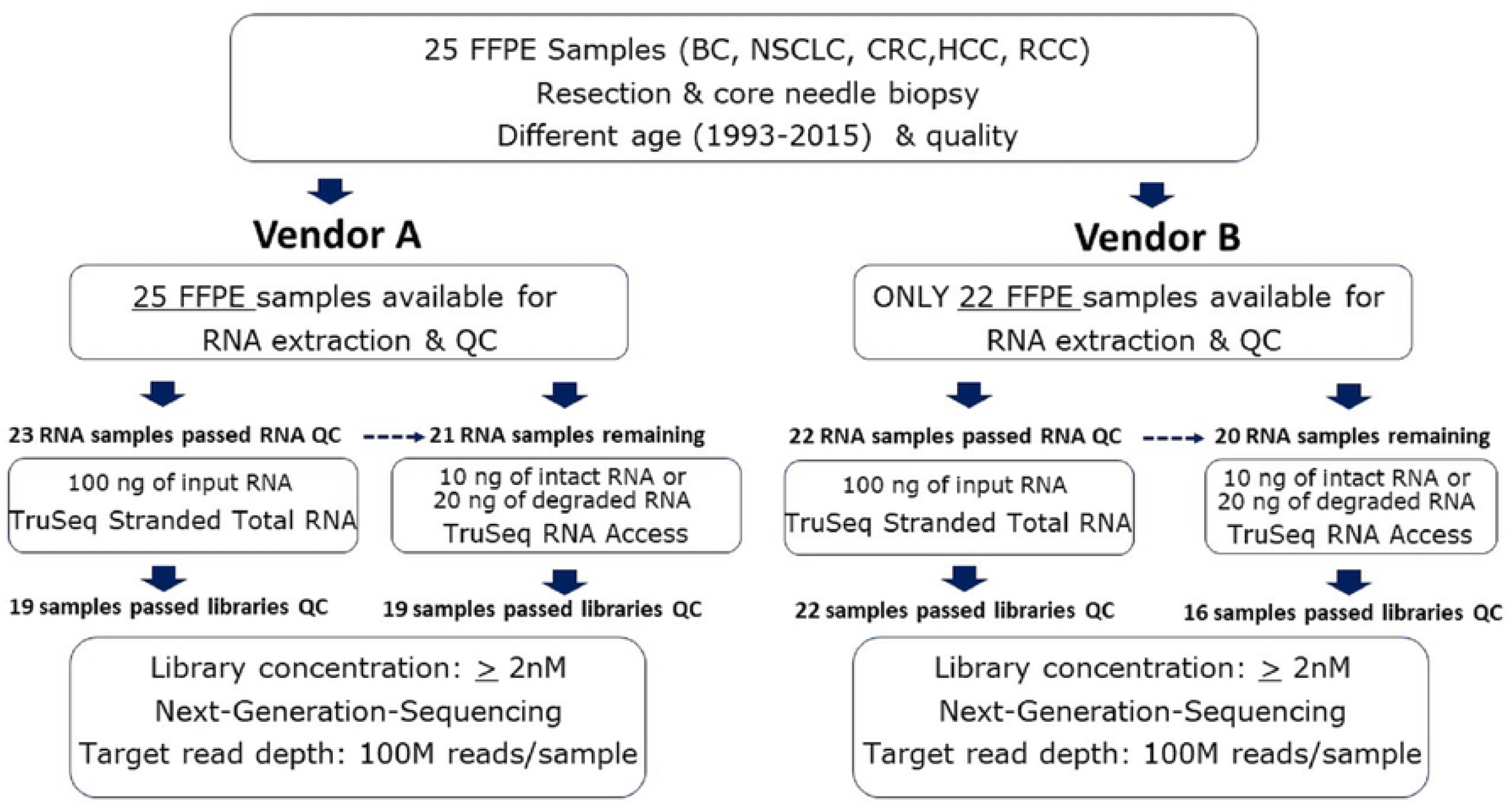
Study design & workflow. Schematic of the sample flow through the two vendors and two protocols. The number of samples processed at each step is noted.

The samples were first sent to vendor A. There, 25 FFPE samples were extracted and analyzed for RNA integrity and quality. A total of 23 FFPE specimens had sufficient yield (>100ng, average DV_200_ is 28% with the range from 5% to 51%) to proceed to TruSeq Stranded Total RNA library preparation. Four sample libraries had a final concentration of less 2nM and therefore did not proceed to sequencing. A total of 19 libraries had sufficient yield to proceed to sequencing. All sequenced samples generated adequate reads (100M or greater). Since more RNA sample is required in the TruSeq Stranded Total RNA protocol, only 21 samples had sufficient RNA remaining (total yield> 20ng, average DV_200_ is 27 with the range from 5 to 51) for the TruSeq RNA Access library preparation kit. Two Access library failed library QC and then 19 samples were sequenced.

Due to insufficient FFPE slides for three samples, Vendor B performed RNA extraction on the 22 remaining FFPE samples. All extracted RNA passed extraction QC (total yield>100 ng, average DV_200_ is 44% with the range from 12% to 72%) to proceed to TruSeq Stranded Total RNA library preparation. All 22 samples had sufficient yield and proceeded to sequencing. The remaining 20 RNA samples (total yield >20ng, average DV_200_ is 44% with the range from 12% to 72%) were re-prepared using the Illumina TruSeq RNA Access library kit and 16 samples passed the library QC for sequencing. Fifteen samples were available from both vendors and both kits and were thus used for further analysis.

For each vendor and kit, Table 1 shows the mean (and range or standard deviation) for process QC measures. The library preparation output is characterized by the average fragment size (measured by Bioanalyzer) and the library concentration. The sequencing output is characterized by the number of reads and a variety of metrics concerning the read alignment rates to exons and ribosomal regions. S1 Table shows the sample annotations and pre-sequencing QC results for each sample.

**Table 1.** Illumina TruSeq RNA Access versus TruSeq Stranded Total RNA: Overall QC and Alignment Stats.

| Step | QC measure | Vendor A |  | Vendor B |  |
| --- | --- | --- | --- | --- | --- |
|  |  | Total Stranded (n=15) | RNA Access (n=15) | Total Stranded (n =15) | RNA Access (n =15) |
| Library Prep | Average Fragment size: Average (Range) | 318 (260-413) | 309.83 (280-326) | 296.63 (268-324) | 324.5 (280-394) |
|  | Concentration (nM): Average (Range) | 57.43 (18.68-108.81) | 51.24 (3.89- 150.63) | 224.57 (6.51 - 507.85) | 205.81 (9.76 - 600.62) |
| Sequencing | Total paired end reads: Average (Range) | 137M (117-157) | 141M (104-169) | 64.7 M (22.5-191) | 184M (114-294) |
| | %Aligned reads rate: Average ± SD | 89.6 ± 10.8 | 95.2 ± 1.0 | 86.3 ± 7.8 | 92.7 ± 1.5 |
| | %Exonic rate: Average ± SD | 18.5 ± 4.7 | 81.0 ± 2.3 | 41.6 ± 13.2 | 84 ± 2.4 |
| | %Intragenic rate: Average ± SD | 83.1 ± 6.8 | 89.1 ± 2.3 | 81.5 ± 17.6 | 92.1 ± 1.9 |
| | %rRNA rate: Average ± SD | 2.0 ± 1.9 | 2.2 ± 1.6 | 9.9 ± 1.83 | 1.8 ± 2.2 |
| | % Correct strand reads rate: Average ± SD | 94.1 ± 3.3 | 95.9 ± 2.3 | 97.5 ± 1.3 | 97.8 ± 1.5 |

### Alignment statistics

Our results showed that the TruSeq RNA Access library preparation protocol produced higher alignment rates at both vendors (means 95% and 93% vs 83% and 78%; Table 1). Compared to the TruSeq Stranded Total RNA protocol, the TruSeq RNA Access protocol showed marked differences in the percent of reads aligned to exons, introns, and intergenic regions. For TruSeq RNA Access the percentages of exonic reads were over 80% across different quality samples at both vendors, reflecting the high efficiency of the exome pull down by the capture approach. The mean exonic percentages were 81% and 84% with the TruSeq RNA Access kit and 17% and 31% with the TruSeq Stranded Total kit.

S2 Table shows per-sample data, including output from Picard, genes detected (>= 1 read), and the 90^th^ percentile gene count.

### Agreement between vendors

The two kits showed similar correlation between vendors. With the TruSeq Stranded Total RNA kit, the average per-sample Spearman correlation between vendors was 0.87 and the average Pearson correlation was 0.76. With the TruSeq RNA Access kit, the average per-sample Spearman correlation between vendors was 0.89 and the average Pearson correlation was 0.73. Across individual samples, the correlation between vendors ranged from R=.94 to R=.01 (Fig 2).

**Figure 2.**
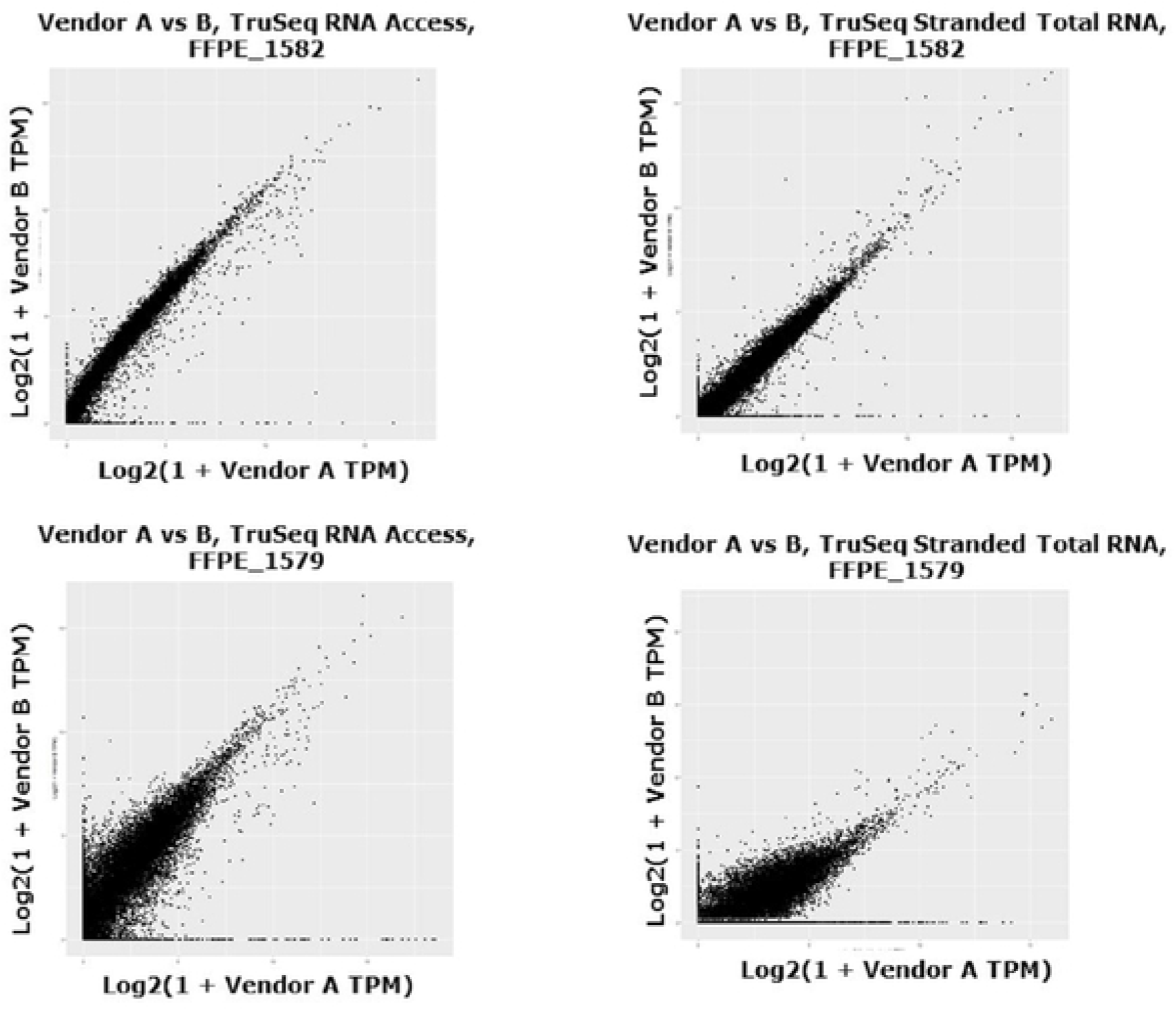
Example cross-vendor scatterplots. The overall correlation between vendors ranged from excellent (eg, A: FFPE_1582 in both TruSeq Stranded Total RNA (R=0.873, rho=0.927) and TruSeq RNA Access kit (R=0.858, rho=0.927)) to moderate (eg, B: FFPE_1579 in both TruSeq Stranded Total RNA (R=0.012, rho=0.760) and TruSeq RNA Access kit (R=0.131, rho=0.869)).

### Agreement between protocols

QC data for FFPE RNA extraction, library preparation, and sequencing from both vendors are comparable. Both vendors achieved similar agreement between protocols. Amongst the 15 samples available in all four datasets, the average Spearman correlation between protocols at vendor A was 0.81 and at vendor B it was 0.83. The average Pearson correlation was 0.13 at vendor A and 0.22 at vendor B.

While the scatterplots for the individual samples (see S1 Fig for the complete set) make it clear that the correlation between protocols is generally good, it is difficult to tell whether there is any systematic difference between protocols. For this, we used Q-Q plots; deviations from a straight line in these plots suggest systematic differences in the dynamics between the two protocols. A number of samples show off-diagonal behavior at the upper end of expression, suggesting that either the TruSeq Stranded Total RNA protocol is saturating or that the TruSeq RNA Access protocol is over-amplifying very highly expressed genes (Fig 3 for one example; see S3 Fig for full set).

**Figure 3.**
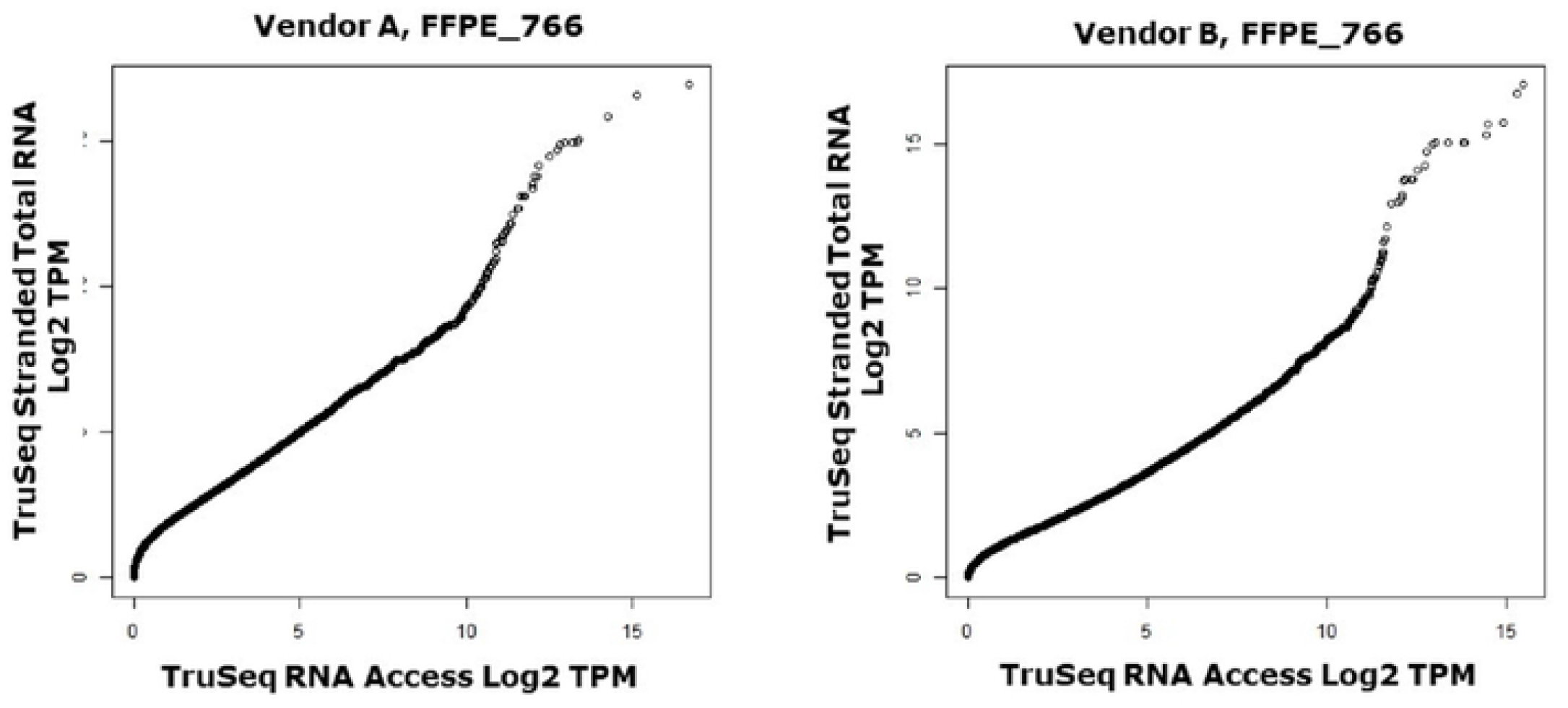
Q-Q plots. The Q-Q plots help visualize the shape of the correlation or distribution between the two kits. Here, the data for sample FFPE_766 is shown from both vendors. At both, the majority of the plot shows a straight diagonal line, indicating identical distribution of TPMs for most percentiles. However, the highest percentiles diverge from the diagonal and the TruSeq Stranded Total RNA kit shows higher levels than the TruSeq RNA Access kit. The plot should not be interpreted to mean that either kit is necessarily correct; only that the highest expressed genes in the TruSeq Stranded Total RNA kit yield higher TPM values than the highest expressed genes in the TruSeq RNA Access kit. The plots also show divergence at very low expression values, potentially genes which are not present in the Access probe set and thus generate no signal in the TruSeq RNA Access results while generating some signal in the TruSeq Stranded Total RNA kit.

We were interested in whether any QC factors (e.g. RNA input, library concentration), especially those obtained before sequencing, might predict the correlation between vendors. Such a predictor could be used in future experiments to distinguish samples likely to produce high quality output from those which may not. We thus examined the Spearman correlation between QC factors (the RNA quantity from each vendor and the library concentration from each vendor) and the Spearman correlation between the data from the two vendors. Table 2 summarizes the results, which suggest that library concentration is better correlated with consistency between vendors than is the RNA quantity. For example, for the TruSeq Stranded Total RNA data, the Spearman correlation of Vendor A’s library concentration with the correlation between Vendor A’s results and Vendor B’s results is 0.96. Fig 4 shows the data in detail, plotting the cross-vendor Spearman correlation vs library concentration.

**Figure 4.**
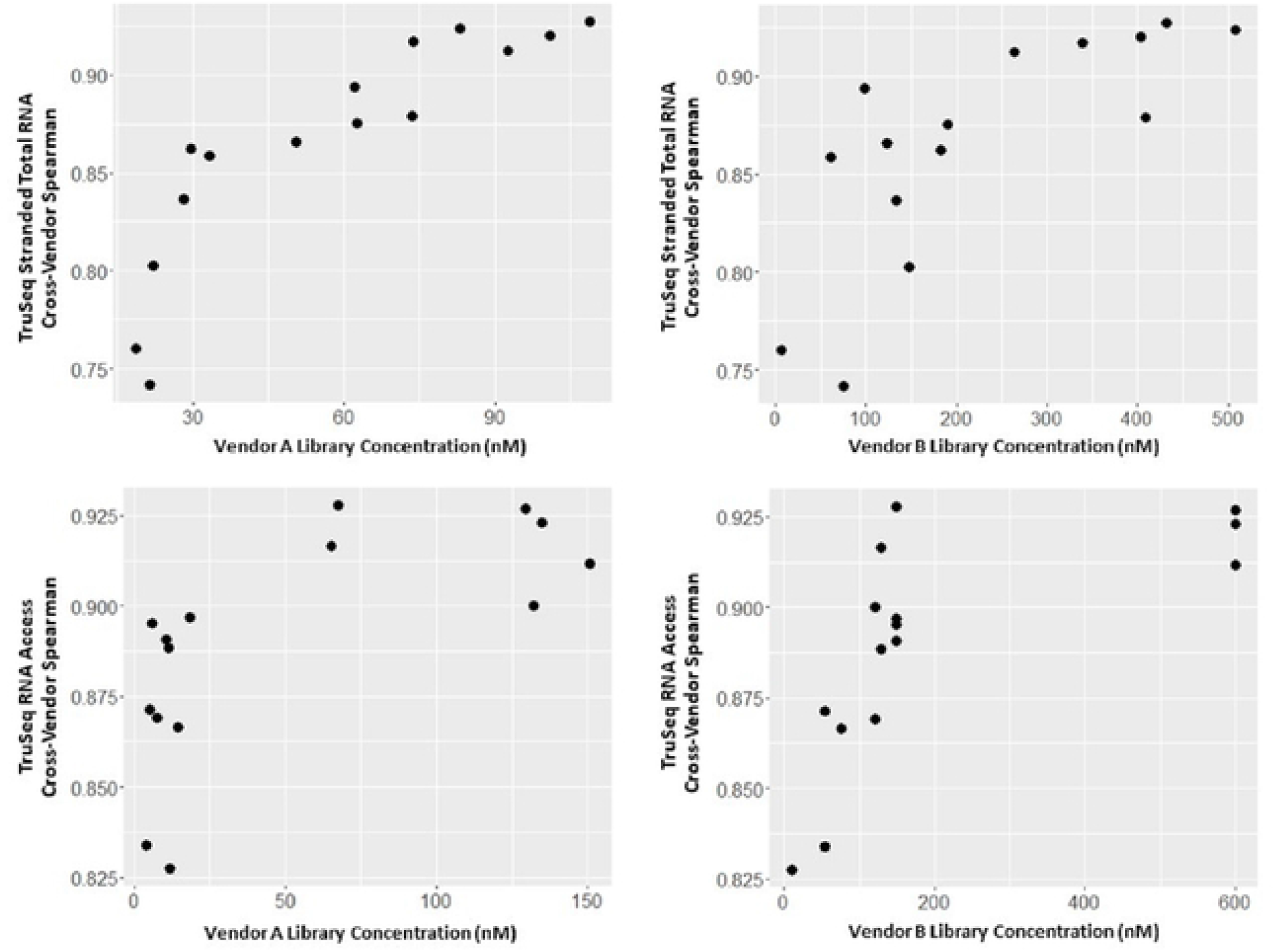
Cross-vendor correlation vs library concentration. Examination of the cross-vendor correlations compared to various common QC statistics suggested that the library concentration was most informative in predicting the cross-vendor correlation. Plotted here are the cross-vendor correlation values vs library concentration. For the TruSeq Stranded Total RNA kit, there is a trend of increasing (though perhaps non-linear) correlation as library concentration increases. For TruSeq RNA Access, there appears to be notably better results from library concentrations above 50nM.

**Table 2.** Spearman correlations between library QC factors (total RNA extracted, library concentration) and the spearman correlation between vendors of the eventual gene-level quantification. This uses cross-vendor correlation as a proxy for the quality of the result and looks at which QC factors might predict that result quality.

|  | TruSeq RNA Access | TruSeq Stranded Total RNA |
| --- | --- | --- |
| Vendor A µg | 0.421 | 0.264 |
| Vendor B µg | 0.481 | 0.35 |
| Vendor A Library Conc | 0.732 | 0.964 |
| Vendor B Library Conc | 0.812 | 0.821 |

## Discussion/Conclusions

RNA-seq is a powerful technology in transcriptome profiling. However, the challenge remains to choose suitable RNA-seq protocols for oncology FFPE specimens with degraded and low quantity RNA sample material. To guide the experimental design of clinical FFPE sample RNA-Seq, we conducted a comparison study using two Illumina library preparation protocols at two vendors for analyzing human RNA isolated from FFPE tissues.

Our results showed that both kits have the similar cross-vendor correlations, suggesting that both protocols offer reproducible results between different operators. However, the two library preparation kits yielded substantial differences in output consistent with the different approaches that the two kits take. Since more RNA sample is required in the Total TruSeq Stranded Total RNA protocol, only 20 samples had remaining RNA for TruSeq RNA Access library preparation. Thus, the TruSeq RNA Access protocol may be the preferred library prep for samples with limited quantity. The Illumina TruSeq RNA Access Library kit generated a higher fraction of reads from protein coding regions compared to other genomic regions; thus, it is a more efficient way to assay the expression of protein coding genes given a limited sequencing budget.

While the TruSeq RNA Access kit may be preferred for difficult samples, the resulting data may not be completely comparable to data from the TruSeq Stranded Total kit or to other kits based on ribosomal depletion and random priming. Further, the two library preparation kits yielded different dynamics of the output transcripts-per-million data at high expression levels where the TruSeq Stranded Total protocol tended to capture genes with higher expression and GC content. The probe-based selection of TruSeq RNA Access libraries may influence output differently than the random priming in other kits. Finally, the probe selection approach precludes certain downstream analyses, such as testing for viral or bacterial content, that may be valuable in some settings. Thus, the lower sequencing costs should be weighed carefully against the anticipated uses for the data to decide which is appropriate for a given experiment. However, since the probe selection step in the RNA Access protocol may bias results compared to other platforms, further exploration may be needed.

In our study, two samples, ages 14 years and 16 years, failed RNA extraction QC in Vendor A, suggesting the influence of age on sample quality on RNAseq library preparation and sequencing. S1 Table lists the detailed reasons for all failures in extraction and library preparation. Interestingly, library concentration appeared to be the best predictor of reproducibility across vendors and thus may be a preferred QC metric for future experiments on FFPE material. While this finding may be useful in avoiding sequencing samples with a low chance of providing quality data, it is not optimal as it can only be applied after the FFPE material is consumed, RNA extracted, and the work of library preparation is completed.

In summary, the quality and quantity of sequencing data obtained through RNA-Seq were strongly influenced by the type of the sequencing library kits. Illumina TruSeq RNA Access library protocol could be a low-cost solution on highly degraded and limited FFPE samples, such as those from clinical studies in which the FFPE quality is severely compromised.

## Acknowledgement

The authors would like express gratitude to Alice Huang for her great support on the conceptualization and reviewing the manuscript. The authors also would like express gratitude to Stefan Pinkert for his great support on sequencing data QC.

## Supporting information

**S1 Table**: Sample characteristics and data from pre-analysis QC (eg, RNA extraction, library preparation).

**S2 Table:** Post-analysis QC output.

**S1 Fig:** Complete set of cross-vendor scatterplots

**S2 Fig:** Full set of Q-Q plots.

